# Microglia and border-associated mouse macrophages maintain their embryonic origin during Alzheimer’s disease

**DOI:** 10.1101/2021.07.05.451201

**Authors:** Xiaoting Wu, Takashi Saito, Takaomi C. Saido, Anna M. Barron, Christiane Ruedl

## Abstract

Brain microglia and border-associated macrophages (BAMs) display distinct spatial, developmental, and phenotypic features. Although at steady-state, the origins of distinct brain macrophages are well-documented, the dynamics of their replenishment in neurodegenerative disorders remain elusive, particularly for disease-associated microglia (DAMs) and BAMs. In this study, we conducted a comprehensive fate-mapping analysis of murine microglia and BAMs and their turnover kinetics during Alzheimer’s disease (AD) progression. We used a novel inducible AD mouse model to investigate the contribution of bone marrow cells to the pool of foetal-derived brain macrophages during the development of AD. We demonstrated that microglia and DAMs remain a remarkably stable embryonic-derived population even during the progression of AD pathology, indicating that neither parenchymal macrophage subpopulation originates from, nor are replenished by, monocytes. At the border-associated brain regions, bona fide CD206^+^ BAMs are minimally replaced by monocytes, and their turnover rates are not accelerated by AD. In contrast, all other myeloid cells are swiftly replenished by bone marrow progenitors. This information further elucidates the turnover kinetics of these cells not only at steady-state, but also in neurodegenerative diseases, which is crucial for identifying potential novel therapeutic targets.

**Impact statement:** Inducible fate-mapping analysis demonstrates that neither microglia, disease-associated microglia nor border-associated macrophages are replenished by bone marrow-derived cells in Alzheimer’s disease.

## Introduction

Microglia are brain parenchymal macrophages that are unique among tissue-resident macrophages due to their primitive yolk sack origin, self-renewal properties (Ajami et al., 2007) and independence from adult hematopoiesis (Ginhoux et al., 2010; Schulz et al., 2012; Sheng et al., 2015). These cells are essential for normal neuronal development, brain function, and central nervous system (CNS) homeostasis and dysfunction is associated with severe brain neuropathologies (Leng and Edison, 2021; Sevenich, 2018). In neurodegenerative diseases, such as Alzheimer’s disease (AD), microglia form a barrier around amyloid plaques, thereby protecting against the neurotoxicity of aggregated beta-amyloid (Aβ) (Condello et al., 2015). Disruption of microglial activity, which is often related to aging and inflammation, can promote this neurotoxicity (Salter and Stevens, 2017) and affect the clearance of Aβ aggregates (Floden and Combs, 2011). The resulting increase in plaque formation (Spangenberg et al., 2019) leads to the progression of pathogenesis.

In several neurodegenerative diseases, the gradual appearance of a subpopulation of disease-associated microglia (DAMs), also known as neurodegenerative microglia (MGnD), correlates with the progression of disease pathology (Deczkowska et al., 2018; Keren-Shaul et al., 2017; Krasemann et al., 2017). In AD, the accumulation of these highly phagocytic cells around the Aβ deposits (Kamphuis et al., 2016; Yin et al., 2017) mitigates Aβ-associated tau seeding and spreading (Gratuze et al., 2021). These DAMs may play a protective role in neurodegeneration, thus implicating targeted manipulation as a therapeutic strategy. Although the origin of the myeloid cells that accumulate around the Aβ deposits is controversial, early studies indicated selective migration of monocytic cells towards the Aβ plaques (Hohsfield and Humpel, 2015), while more recent research suggests that these cells are derived from resident embryonic-derived microglia (Reed-Geaghan et al., 2020; Shukla et al., 2019).

In addition to the macrophages in brain parenchyma, microglia-independent macrophages are also found at the border regions, such as the subdural meninges (SDM), including the pia and arachnoid mater, the dura matter (DM), and the choroid plexus (CP) (Brioschi et al., 2020; Utz et al., 2020). Single-cell RNAseq analysis revealed that these border-associated macrophages (BAMs) represent a family of different subpopulations (Van Hove et al., 2019). However, the contributions of BAMs to CNS integrity and neurodegeneration remain to be elucidated.

To investigate the influence of AD on the macrophage CNS landscape, we analyzed the phenotype and turnover kinetics of different myeloid cells in the brain parenchyma and border-associated tissues. We used a novel AD mouse model generated by back-crossing *App*^NL-G-F^ knock-in (APP-KI) mice (Saito et al., 2014) with a *Kit*^MerCreMer^/*R26*^YFP^ fate-mapping mouse strain (Sheng et al., 2015) to monitor bone marrow (BM)-driven replenishment of brain myeloid cells during disease progression. We also investigated the origins of DAMs since the ontogeny of “activated” microglia in AD is still a matter of debate.

## Results and discussion

In this study, we conducted a comprehensive fate-mapping analysis of murine brain microglia and BAMs and their turnover kinetics during the progression of AD.

Mouse models of AD have been instrumental in clarifying the cellular and molecular mechanisms underlying this irreversible brain disorder. Transgenic mice overexpressing proteins linked to familial AD (5xFAD), single mutant amyloid precursors (APP), or double mutant APP and presenilin (APP-PS1) have been used in many studies (Sasaguri et al., 2017). In our study, we exploited an APP-KI mouse strain expressing a mutant form of humanized APP, the parent protein of Aβ, knocked-in under the control of the endogenous promoter. This model avoids some disadvantages associated with transgenic APP models, including artefacts caused by overexpression of other APP fragments in addition to Aβ, non-physiological cell-type expression and potential insertion site disruption (Saito et al., 2014). The APP-KI mouse line expresses three human AD-associated mutations that promote the progressive accumulation of Aβ through its increased production of Aβ, particularly the more toxic Aβ42 form, as well as increasing Aβ aggregation and reducing degradation. APP-KI mice develop progressive Aβ accumulation from 2 months of age, thus mimicking several aspects of human AD, including microgliosis and synaptic loss (Fig. 1- figure supplement 1A).

First, we characterized the myeloid cell landscape in different brain regions of healthy young mice (8 weeks) as well as aged WT and APP-KI mice (12 months). In our multiparameter flow cytometry and uniform manifold approximation and projection (UMAP) analyses, we included a panel of myeloid markers to delineate microglia, DAMs, monocyte-derived macrophages (MdCs), border-resident macrophages (BAMs), neutrophils, monocytes and, eosinophils.

P2RY12^+^ microglia were the main CD45^int^ F4/80^hi^ cell population in the brain parenchyma of all mice. AD mice showed a substantial increase in the frequency of P2RY12^low^CD11c^+^ DAMs, which were absent in the brain of young mice and constituted only a minor fraction in aged mice (Fig. 1 A-C). In accordance with reports of the appearance of DAMs during neurodegeneration in other AD transgenic mouse models, such as 5xFAD and APP/PS1 (Keren-Shaul et al., 2017; Mrdjen et al., 2018), this phenotype was recapitulated in APP-KI mice in our study (Fig. 1- figure supplement 1B and C). The remaining myeloid cell subsets detected in the brain parenchyma, such as neutrophils (Ly6G^+^), monocytes (Ly6C^hi^ and Ly6C^int^), monocyte-derived macrophages (MdCs) (F4/80^int^MHCII^hi^ and F4/80^int^MHCII^−^CD11a^+^), eosinophils (Siglec-F^+^), and cDCs (CD11c^hi^MHCII^hi^), were mainly restricted to the CD45^hi^ gate. Surprisingly, we did not observe an enhanced infiltration of blood-derived inflammatory cells in the brain during disease progression. Even in aged APP-KI mice, the infiltration by Ly6C^hi^ monocytes and Ly6G^+^ neutrophils was comparable to that in age-matched controls (Fig. 1 A-C).

**Fig. 1:**
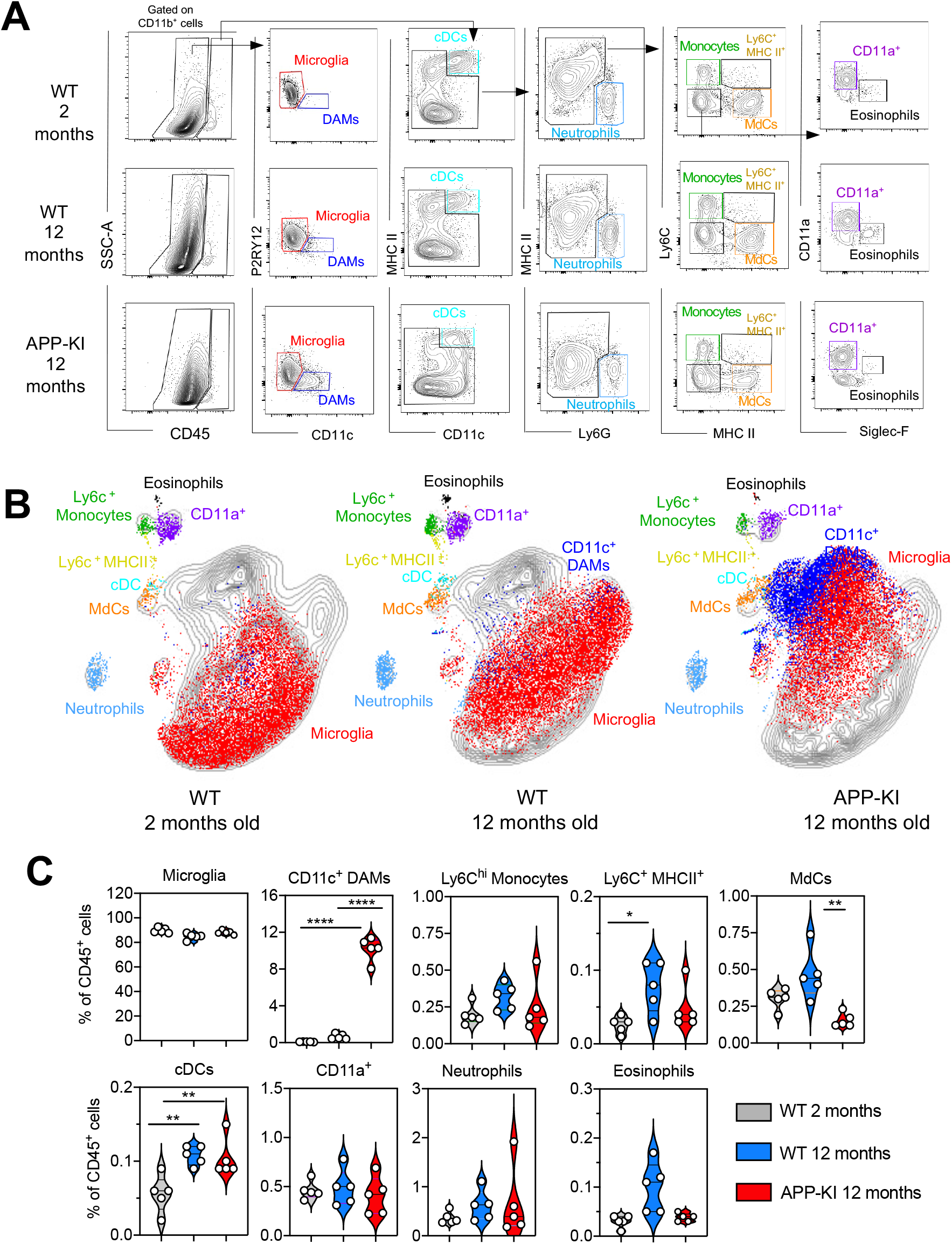
Myeloid cell profiling in healthy and AD parenchymal brains. (A) Representative gating strategy used to visualize distinct myeloid cell subpopulations in young WT (aged 8 weeks), WT (aged 12 months) and diseased APP-KI (aged 12 months) mice. Microglia: CD45^int^F4/80^hi^P2RY12^+^CD11c^−^; DAMs: CD45^int^F4/80^hi^P2RY12^+^CD11c^+^; cDCs: CD45^hi^CD11c^hi^MHCII^+^; neutrophils: CD45^hi^CD11b^hi^Ly6G^+^; monocytes: CD45^hi^F4/80^int^CD11b^int^Ly6C^+^MHCII^−^; monocyte-derived macrophages (MdCs): CD45^hi^F4/80^int^CD11b^int^Ly6C^−^MHCII^+^; CD11a^+^ cells: CD45^hi^F4/80^int^CD11b^int^Ly6C^−^MHCII^−^CD11a^+^ and eosinophils: CD45^hi^F4/80^int^CD11b^int^Siglec-F^+^. (B) UMAP analysis displaying 30,000 randomly sampled cells from young WT, aged WT and APP-KI mouse brains analyzed by multicolor flow cytometry (n = 3 mice/group). (C) Violin plots with individual dots illustrating the frequency of different myeloid cell populations within the total parenchymal brain CD45^+^ cell population. Each dot represents the percentage of cells obtained from one brain (n = 5 mice/group). Young mice: grey, aged mice: blue and APP-KI mice: red. Samples were analyzed by two-way ANOVA. \**P* < 0.05; \*\**P* < 0.01; \*\*\*\**P* < 0.0001. For clarity, non-significant values are not shown.

In all three separately analyzed CNS border-associated tissues, a predominant F4/80^hi^CD206^+^ BAM fraction could be further separated into MHCII^+^ and MHCII^low^ subpopulations, which is consistent with previous reports (Van Hove et al., 2019) (Fig. 2 A-C, Fig. 2- figure supplement 1 C). A large population of CD206^+^MHCII^low^ BAMs was preferentially localized in the SDM, whereas these cells were less abundant in the DM and CP. Interestingly, a significant transition from CD206^+^MHCII^low^ to CD206^+^MHCII^+^ BAMs was observed with aging and AD progression. All other myeloid cell populations, including neutrophils, monocytes, MdCs, CD11a^+^ cells, eosinophils, and cDCs, were present in each tissue, although at different frequencies (Fig. 2 A-C, Fig. 2- figure supplement 1 C). Similar to the brain parenchyma, monocyte and neutrophil frequencies were not augmented in any of the three border regions of the aged WT and APP-KI mice, which excludes the possibility that an enhanced inflammatory cell infiltration is caused by aging or neurodegenerative disease progression (Fig. 2 A-C, Fig. 2- figure supplement 1 C).

**Fig. 2:**
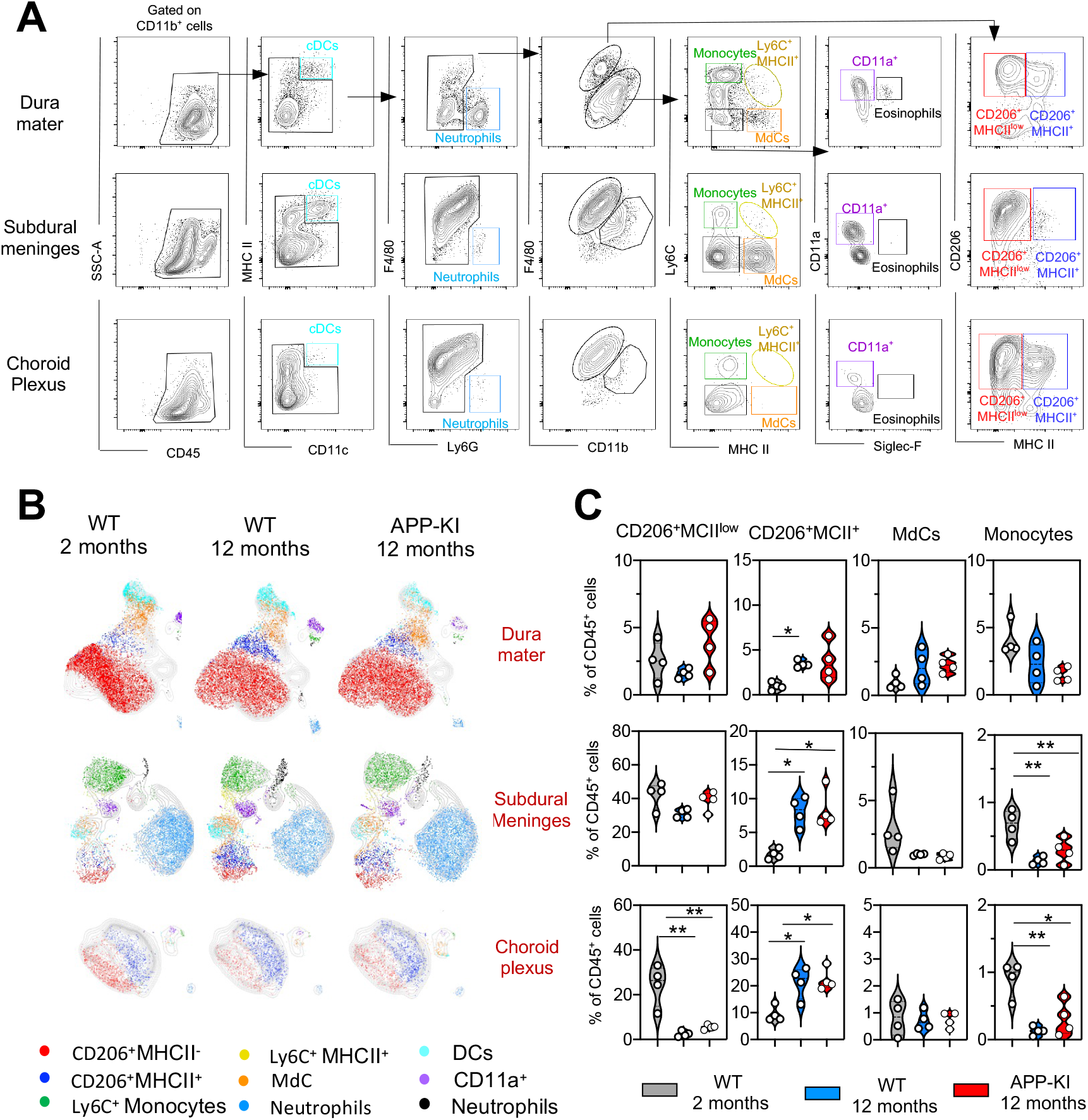
Characterization of myeloid cells in distinct brain border regions in health and AD. (A) Representative gating strategy used to visualize distinct myeloid cell subpopulations in the dura matter (DM), subdural meninges (SDM) and choroid plexus (CP). Young WT mice (aged 8 weeks) were analyzed. Cell classification as shown in the legend for Figure 1. (B) UMAP analysis displaying 8500 (DM), 13000 (SDM), 3700 (CP) randomly sampled cells from young WT, aged WT and APP-KI DM, SDM and CP analyzed by multicolor flow cytometry (n = 9 mice). (C) Violin plots with individual dots illustrating the frequency of different myeloid cell populations within the total non-parenchymal CD45^+^ cell population of the DM, SDM and CP. Young mice: grey, aged mice: blue and APP-KI mice: red. Each dot represents the percentage of cells obtained from pooled border regions (n = 2–3 pooled mice). Samples were analyzed by two-way ANOVA. \**P* < 0.05; \*\**P* < 0.01. For clarity, non-significant values are not shown.

To investigate the origins and replenishment kinetics of distinct brain macrophage subpopulations, we crossed the APP-KI mouse line with the Kit^MerCreMer^/*R26*^YFP^ fate-mapping mouse strain. The resulting APP-KI-Kit^MerCreMer^/*R26*^YFP^ inducible adult fate-mapping mouse model allowed us to irreversibly label Kit-expressing BM-progenitors and trace them in different brain regions during the progression of AD. To induce the YFP label in Kit-expressing BM progenitor cells, *Kit*^MerCreMer^/*R26*^YFP^/APP-KI mice and the corresponding *Kit*^MerCreMer^/*R26*^YFP^ controls were administered TAM at 2, 4, 6 and 8 months of age. Parenchymal and non-parenchymal brain cells were isolated separately and analyzed by multicolor flow cytometry.

In the parenchyma, a strong YFP signal was limited to CD45^hi^ cells and was barely detectable in the CD45^int^ fraction, which included the microglia and DAMs, in both WT and AD mice (Fig. 3 A-C). In fact, microglia showed minimal YFP-labeling, which further confirmed their embryonic origin and BM-independence (Sheng et al., 2015). During AD progression, microglial YFP-labeling remained minimal, indicating that the ongoing neurodegeneration did not promote their replenishment by BM-derived cells. Similarly, DAMs, which were increasingly formed during AD progression (Fig. 1- figure supplement 1A), maintained a low YFP-labeling profile, suggesting that despite their phenotypic shift, these cells preserved their embryonic signature and were not replaced by BM-derived cells during evolution of the disease (Fig. 3). To verify whether DAMs retained their fetal lineage, we performed E7.5 embryonic labeling of *Kit*^MerCreMer^/*R26*^YFP^/APP-KI mice and analyzed microglia and DAMs obtained from disease-affected offspring aged 9 months. Despite the AD-associated pathology, we not only confirmed that DAMs maintained their embryonic origin, but showed that microglia and activated DAMs share the same yolk sac origins (Fig. 3 D). The kinetics of monocyte replacement were extremely rapid, with all monocytes YFP-labeled 2 months after induction, indicating that these cells are exclusively BM-derived (Fig. 3 B, lower panel, left). In contrast, fetal-derived MdCs were gradually replaced by BM-derived cells and were fully replenished at 8 months post-induction, with no significant difference between WT and AD mice (Fig. 3 B, lower panel, right).

**Fig. 3:**
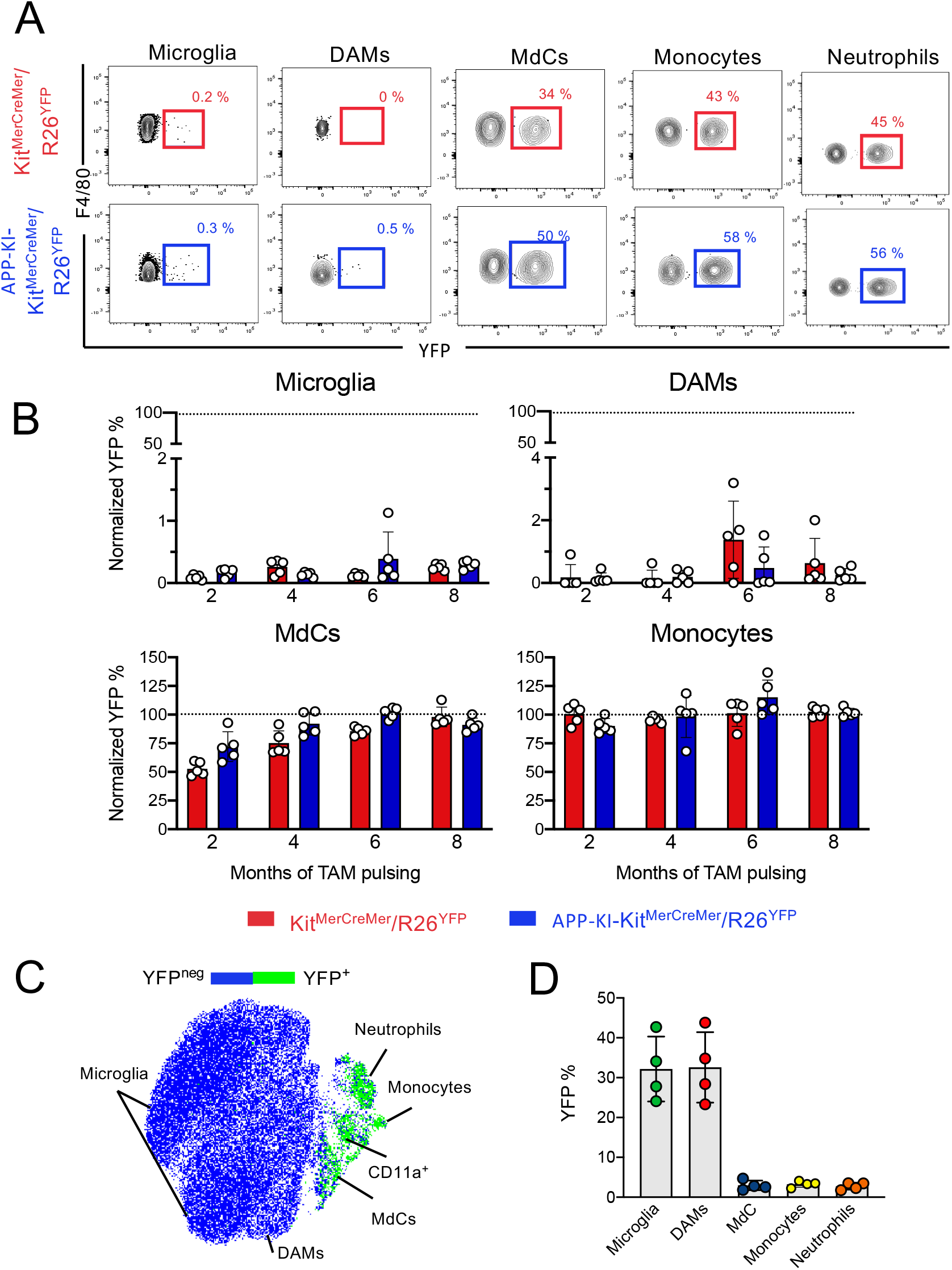
Microglia and disease-associated microglia maintain their embryonic origin during Alzheimer’s disease. (A) Representative flow cytometry contour plots indicating the YFP-labeling of parenchymal microglia, DAMs, MdCs, monocytes and neutrophils. (B) *Ki*t^MerCreMer^/*R26*^YFP^ and APP-KI-*Ki*t^MerCreMer^/*R26*^YFP^ mice (aged 2, 4, 6 and 8 months) received 4 mg tamoxifen (TAM) by oral gavage for five consecutive days and were sacrificed at 10 months of age. Bar charts with individual data points showing the percentage of YFP^+^ parenchymal microglia, DAMs, MdCs and monocytes obtained from *Ki*t^MerCreMer^/*R26*^YFP^ (red bars) and APP-KI-*Ki*t^MerCreMer^/*R26*^YFP^ mice (blue bars) after normalization to the percentage of YFP^+^ neutrophils. Data represent the mean ± SD (n = 5 mice). (C) UMAP representation showing the YFP-labeled cell populations in green and the YFP-negative fraction in blue. (D) Embryonic fate-mapping. A single pulse of tamoxifen was administered to APP-KI-Kit^MercreMer^/R26 pregnant mice at E7.5. Offspring were analyzed at 9 months of age. Bar charts with individual data show the percentages of YFP-labeled parenchymal microglia, DAMs, MdCs, monocytes and neutrophils. Data represent the mean ± SD (n = 4 mice). For each time-point (2, 4, 6 and 8 months), the significance of differences between the WT and AD groups was analyzed by Student’s *t*-test (two-tailed). For clarity, non-significant values are not shown.

Using our fate-mapping mouse model, we then analyzed myeloid cells residing in distinct CNS border-associated regions. In adult fate-mapping, both CD206^+^ BAM subpopulations showed weaker YFP-labeling compared to MdCs, which were already fully replenished within 2 months in all three regions (Fig. 4). In the DM, higher YFP-labeling was significantly higher in CD206^+^MHCII^+^ BAMs than that in CD206^+^MHCII^low^ BAMs (Fig. 4, upper panel). Dural CD206^+^MHCII^+^ BAMs were characterized by a slow, but consistent BM-cell replacement over time, reaching approximately 35% YFP-positivity by 8 months post-induction. In contrast, the turnover rate of both BAM populations located in the SDM was markedly slower (Fig. 4, middle panel), which suggests that their niche is less accessible to BM-dependent replenishment. With its fenestrated blood vessels, the DM has a greater capacity to support peripheral cell traffic. In contrast, the strong tight junctions of the blood vessels in the SDM limit cell exchange with the periphery (Mastorakos and McGavern, 2019). Previous studies have shown a rapid BAM turnover in the CP supported by partial input from the circulation (Goldmann et al., 2016; Van Hove et al., 2019). However, although both MHCII^low^ and MHCII^+^CD206^+^ BAMs displayed a dual origin in our fate-mapping mouse model, their replenishment from BM-derived cells was extremely slow, with the YFP signal reaching only 10%–15% by 8 months after induction. More than 85% of CP BAMs retained their embryonic phenotype (Fig. 4, lower panel). The lack of a significant difference in the YFP-labeling profiles of BAMs from healthy or AD mice indicates that the development of AD neither supports nor accelerates BAM replacement by BM-progenitors (Fig. 4). On the other hand, non-parenchymal monocytes (data not shown) and MdCs were swiftly replaced by BM cells, with 100% YFP-positivity at 2 months post-induction (Fig. 4, right).

**Fig. 4:**
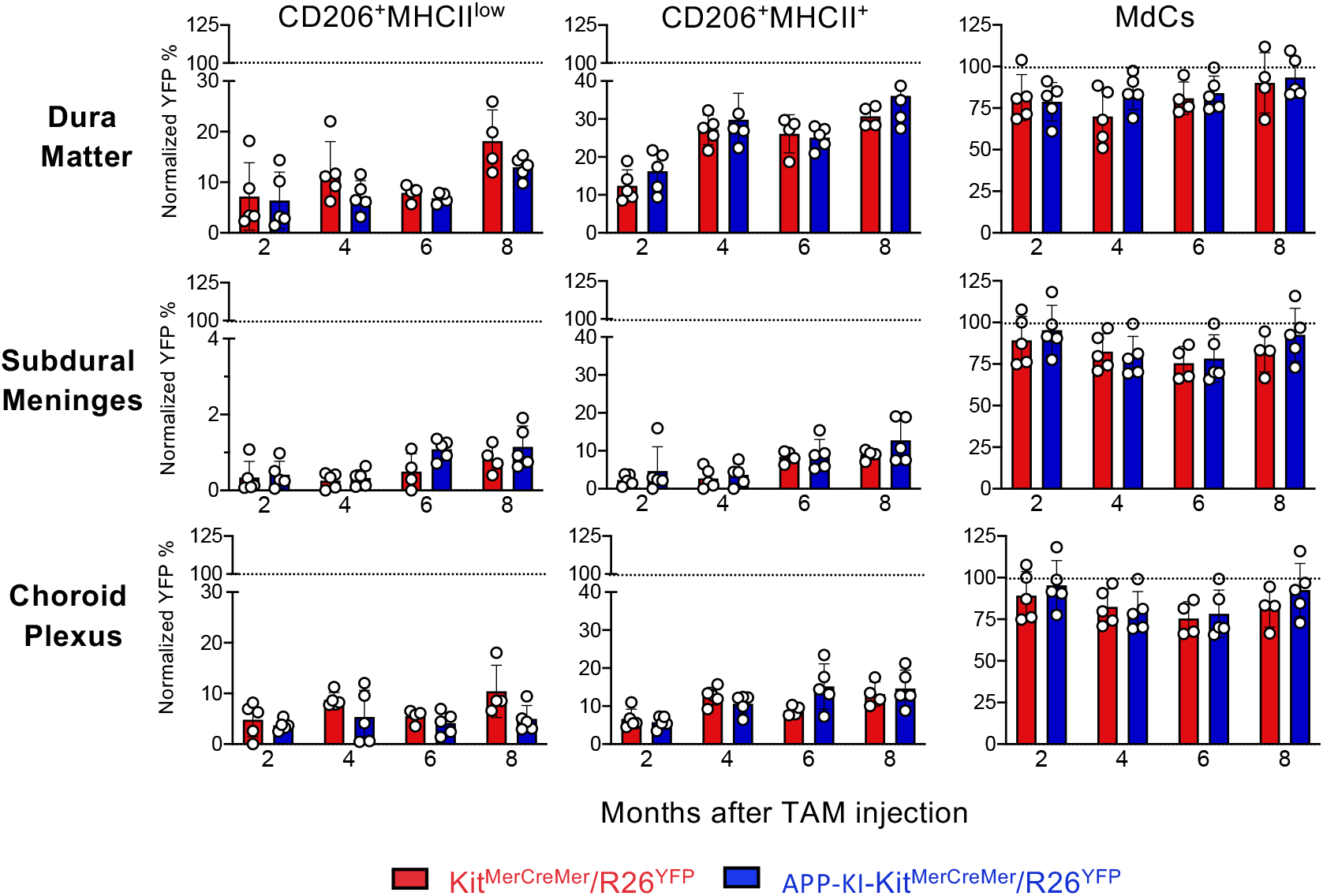
AD does not accelerate the turnover kinetics of border-associated macrophages. Mice were treated and analyzed as described in legend for Figure 3. Bar charts with individual data points showing the percentages of YFP^+^ non-parenchymal BAMs (CD206^+^MHCII^low^ and CD206^+^MHCII^low^ cells) and MdCs obtained from *Ki*t^MerCreMer^/*R26*^YFP^ (red bars) and APP-KI-*Ki*t^MerCreMer^/*R26*^YFP^ mice (blue bars) after normalization to the percentage of YFP^+^ neutrophils. Data represent the mean ± SD (n = 4 samples of 2–3 pooled mice). For each time-point (2, 4, 6 and 8 months), the significance of differences between the WT and AD groups was analyzed by Student’s *t*-test (two-tailed). For clarity, non-significant values are not shown.

The contribution of monocytes to the pool of microglia is unclear. The monocyte-to-microglia transition occurs in the brain, but only under certain conditions of inflammation or injury, such as meningitis (Djukic et al., 2006) and neonatal stroke (Chen et al., 2020). However, the original yolk sac embryonic microglia identity is preserved during experimental autoimmune encephalomyelitis (Ajami et al., 2011; Jordao et al., 2019), although infiltrating monocytes are recruited to the brain during the progression of this neuroinflammatory disease. In an elegant parabiosis experiment using two different mouse models of AD (APPPS1-21 and 5xFAD), it was shown that peripheral monocytes do not contribute to the microglial pool surrounding the Aβ plaques (Wang et al., 2016). Here, we extended our analysis and demonstrated that, in the steady-state, microglia, DAMs and BAMs are a remarkably stable embryonic-derived population and their turnover remains unaffected in AD; the progression of this neurodegenerative disease does not promote or sustain their replacement by BM-derived progenitors. Furthermore, we confirmed that the transformation from homeostatic microglia to activated DAMs is not mediated by infiltrating inflammatory monocytes, but imprinted by the local brain microenvironment that is severely perturbed by the AD pathology. This new understanding will be valuable in manipulating the transition of microglia to DAMs as a therapeutic strategy in AD.

## Material and Methods

### Key resources table

| Reagent type or resource | Designation | Source or reference | Identifiers | Additional information |
| --- | --- | --- | --- | --- |
| **Antibody** | Rat anti-mouse CD45 (HI30) | BD Bioscience | Cat#: 748719; RRID: AB_2873123 | FACS (1:600; 100 µl per test) |
| **Antibody** | Rat anti-mouse Siglec-F (clone: E50-2440) | BD Bioscience | Cat#: 746668; RRID: AB_2743940 | FACS (1:600; 100 µl per test) |
| **Antibody** | Rat anti-mouse CD11b (M1/70) | BioLegend | Cat#: 101226; RRID: AB_830642 | FACS (1:600; 100 µl per test) |
| **Antibody** | Rat anti-mouse CD11b (M1/70) | BD Bioscience | Cat#: 565976 ; RRID: AB_2738276 | FACS (1:600; 100 µl per test) |
| **Antibody** | Rat anti-mouse F4/80 (BM8) | Biolegend | Cat#: 123110; RRID: AB_893486 | FACS (1:600; 100 µl per test) |
| **Antibody** | Rat anti-mouse F4/80 (BM8) | BioLegend | Cat#: 123114; RRID: AB_893490 | FACS (1:600; 100 µl per test) |
| **Antibody** | Rat anti-mouse Ly6c (HK1.4) | BioLegend | Cat#: 128030; RRID: AB_2562617 | FACS (1:600; 100 µl per test) |
| **Antibody** | Rat anti-mouse I-A/I-E Antibody (M5/114.15.2) | BioLegend | Cat#: 107636; RRID: AB_2734168 | FACS (1:1,000; 100 µl per test) |
| **Antibody** | Hamster anti-mouse CD11c (N418) | Biolegend | Cat#: 117318; RRID: AB_493568 | FACS (1:600; 100 µl per test) |
| **Antibody** | Hamster anti-mouse CD11c (N418) | BioLegend | Cat#: 117336; RRID: AB_2565268 | FACS (1:600; 100 µl per test) |
| **Antibody** | Rat anti-mouse P2RY12 (S16007D) | BioLegend | Cat#: 848006; RRID: AB_2721469 | FACS (1:600; 100 µl per test) |
| **Antibody** | Rat anti-mouse CD11a (M17/4) | Invitrogen | Cat#: 48-0111-82; RRID: AB_11064445 | FACS (1:600; 100 µl per test) |
| **Antibody** | Mouse anti-mouse CD206 (clone: C068C2) | BioLegend | Cat#: 141734; RRID: AB_2629637 | FACS (1:600; 100 µl per test) |
| **Antibody** | Mouse anti-mouse Ly6G (1A8) | BioLegend | Cat#: 127606; RRID: AB_1236494 | FACS (1:600; 100 µl per test) |
| **Antibody** | Fc-R block (2.4G2) | Self-made | N/A | Blocking step (1:100; 1,000 µl per sample) |
| Chemical compound, drug | DAPI | ThermoFisher Scientific | Cat#: D1306 | (1:1,000) |
| Chemical compound, drug | Collagenase D | Roche | Cat#: 11088882001 | 1 mg/ml |
| Chemical compound, drug | Dispase II | Gibco | Cat#: 17105041 | 2 U/ml |
| Chemical compound, drug | DNase | Roche | Cat#: 04536282001 | 2 U/ml |
| Chemical Compound, drug | Percoll | Merck | Cat#: P4937-500ML |  |
| Chemical compound, drug | Progesterone | Sigma–Aldrich | Cat#: P0130 | 1 mg/mouse |
| Chemical compound, drug | Tamoxifen | Sigma–Aldrich | Cat#: T5648 | For adult labeling, 4 mg tamoxifen for 5 consecutive days by oral gavage; For embryo labeling, pregnant mice (E7.5) were injected via oral gavage once with 2 mg TAM |
| Chemical compound, drug | IMDM | ThermoFisher Scientific | Cat#: 12440046 |  |
| Software, algorithm | FlowJo | TreeStar | FlowJo 10.6<br>RRID: SCR_008520 |  |
| Software, algorithm | GraphPad Prism | GraphPad Software | GraphPad 9.0<br>RRID: SCR_002798 |  |
| Strain, strain background (mouse) | APP-KI<sup>NL-G-F</sup> | T.Saito, T.C. Saido<br>Japan | Saito et al., 2014 |  |
| Strain, strain background (mouse) | *Kit*<sup>MerCreMer</sup>/*Rosa26-LSL-eYFP* (called *Kit*<sup>MerCreMer</sup>/*R26*<sup>YFP</sup>) | Nanyang Technological University, Singapore | Sheng et al., 2015 |  |
| Strain, strain background (mouse) | APP-KI<br>*Kit*<sup>MerCreMer</sup>/*R26*<sup>YFP</sup>) | Nanyang Technological University, Singapore | Described here |  |

### Mice

Fate-mapping *Kit*^MerCreMer^/*R26*^YFP^ mice were generated as previously described (Sheng et al., 2015). The *Kit*^MerCreMer^/*R26*^YFP^ fate-mapping mouse line was crossed with the AD mouse model (APP^NL-G-F^), a knock-in mouse line that co-expresses the Swedish (KM670/671NL), Beyreuther/Iberian (I716F), and Arctic (E693G) mutations and mimics AD-associated pathologies, including amyloid plaques, synaptic loss, and microgliosis as well as astrocytosis (Saito et al., 2014); the generated mouse line was designated APP-KI-*Kit*^MerCreMer^/*R26*^YFP^. Mice were bred and maintained in a specific pathogen-free animal facility at the Nanyang Technological University (Singapore). All animal studies were carried out according to the recommendations of the National Advisory Committee for Laboratory Animal Research and ARF SBS/NIE 18081, and 19093 protocols were approved by the Institutional Animal Care and Use Committee of the Nanyang Technological University.

### Tamoxifen-inducible embryonic and adult fate-mapping mouse model

*Kit*^MerCreMer^/*R26*^YFP^ and APP-KI-*Kit*^MerCreMer^/*R26*^YFP^ fate-mapping mice were used to determine the turnover rates of distinct brain macrophages under normal steady-state or diseased conditions. For adult labeling, a total of 4 mg tamoxifen (TAM) (Sigma–Aldrich, St. Louis, MO, USA) per mouse was administered for five consecutive days by oral gavage as previously described (Sheng et al., 2015). Mice were sacrificed at different time-points for brain collection, subsequent cell isolation, and multiparameter flow cytometric cell analysis. For embryonic labeling, *Kit*^MerCreMer^/*R26*^YFP^ and APP-KI-*Kit*^MerCreMer^/*R26*^YFP^ fate-mapping mice were mated overnight and separated early the next morning. Pregnant mice (E 7.5) received one dose of 2 mg TAM with 1 mg progesterone via oral gavage.

### Isolation of microglia and border-associated macrophages

Brains were removed and DM, SDM and CP were carefully separated from the brain parenchyma. For the DM isolation, the dorsal part of the skull was removed and the dura were peeled away from the skull cap and placed in 2% fetal bovine serum (FBS) in Iscove’s Modified Dulbecco’s Medium (IMDM). The SDM were micro-dissected using micro suture forceps and placed on ice-cold 2% FBS IMDM. For collection of the CP, the ventricles were exposed and the CP was carefully micro-dissected from the lateral ventricles and placed on 2% FBS IMDM.

All tissues were cut into small pieces, which were subsequently incubated with digestion buffer (IMDM supplemented with 2% FBS, 1 mg/mL Collagenase D [Roche], 2 U/mL DNase I [Life Technologies] and Dispase II [Roche]). The digested brain parenchyma, DM, SDM and CP were separately homogenized with a syringe and the resulting homogenous cell suspensions were filtered through a 40-μm cell strainer. Only the parenchyma cells were resuspended in a 40% Percoll (GE Healthcare Life Sciences) and centrifuged at 700 *xg* for 10 min. All obtained cell pellets were resuspended in 0.89% NH_4_CL lysis buffer for 5 min at room temperature to remove contaminating red blood cells. After centrifugation at 350 x*g* for 5 min, the supernatant was discarded and cell pellets were collected for flow cytometry staining.

### Flow cytometry staining

Isolated cells were pre-incubated with 10 μg/ml anti-Fc receptor antibody (2.4G2) on ice for 20 min. Subsequently, cells were stained with different antibodies for 20 min on ice. After washing, cells were further stained with DAPI to exclude dead cells. Finally, cells were washed and resuspended in PBS/2% FBS for analysis on a five-laser flow cytometer (FACSymphony A3 Cell Analyser, BD Bioscience, San Jose, CA, USA). Data were analyzed with FlowJo software (TreeStar, Ashland, OR, USA).

### Statistical analysis

Statistical analysis was performed using GraphPad Prism 9.0.1 software (GraphPad Software, La Jolla, CA, USA). All values were expressed as the mean ± standard deviation (SD) as indicated in the figure legends. Samples were analyzed by Student’s *t*-test (two-tailed) or two-way ANOVA. A *P*-value <0.05 was considered to indicate statistical significance. The number of animals used per group is indicated in the figure legends as “n.”

### Ethics

All animal studies were carried out according to the recommendations of the National Advisory Committee for Laboratory Animal Research and ARF SBS/NIE 18081, and 19093 protocols were approved by the Institutional Animal Care and Use Committee of the Nanyang Technological University.

## Data availability

The original flow cytometry data have been deposited in the NTU Open Access Data Repository (DR-NTU).

## Author contribution

Conceptualization: C.R.; Methodology: X.W. and A.B.; Animal model: T.S. and T.C.S.: Formal analysis: X.W. and C.R.; Writing: C.R.; Visualization: Q.C. and C.R.; Supervision: C.R.; Funding acquisition: C.R.

## Acknowledgments

The authors would like to thank Insight Editing London for proofreading the manuscript before submission. This work was supported by a Ministry of Education Tier 1 grant awarded to C.R.

---

**Fig. 1- figure supplement 1:**
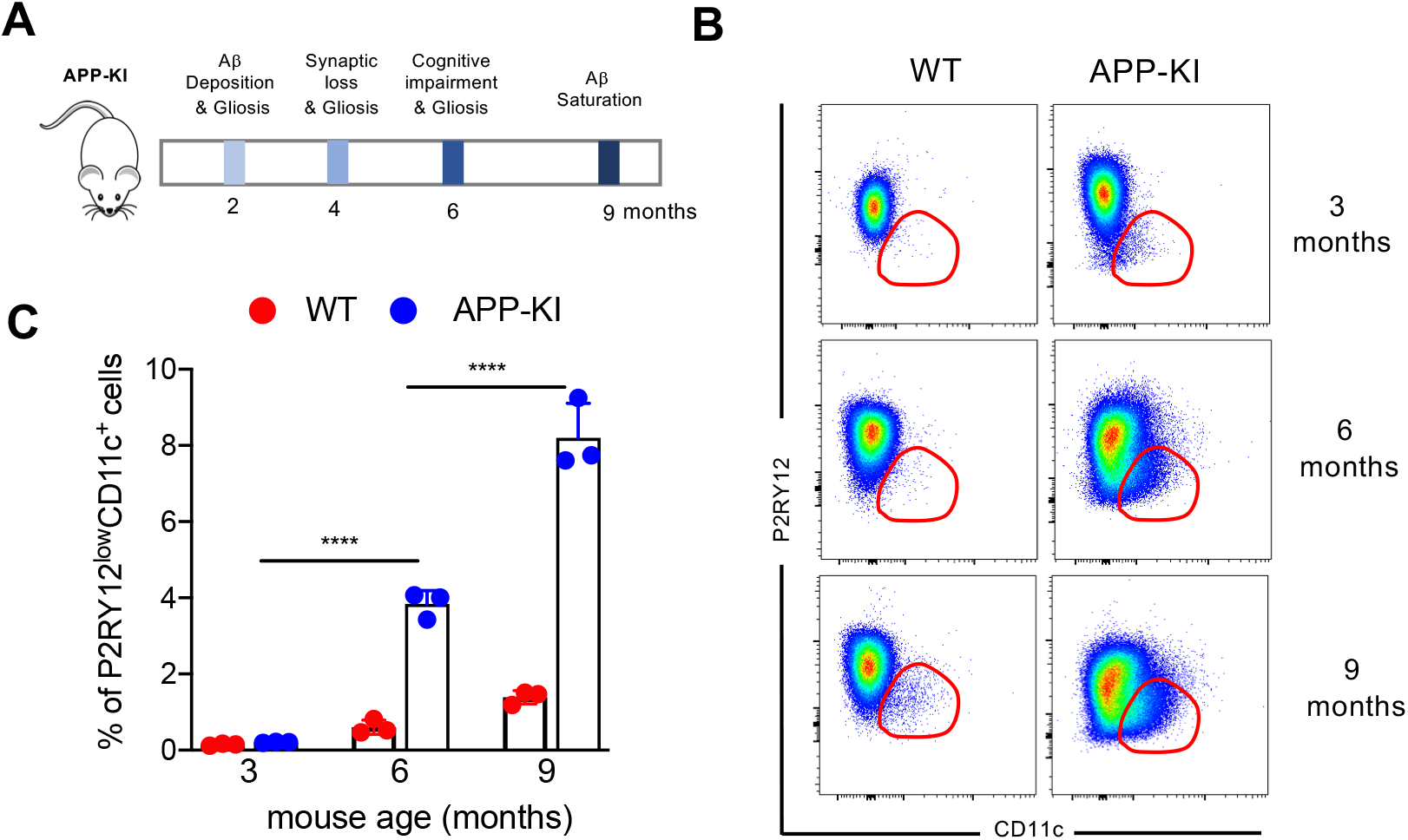
(A) Schematic representation of distinct stages during the progression of AD in the experimental APP-KI^NL-G-F^ transgenic mouse model. (B) Representative flow cytometric analysis showing disease-associated microglia at different stages of AD progression (3, 6 and 9 months) in WT and APP-KI mice. Dot plots showing CD11c on the x-axis and PR2Y12 on the y-axis. (C) Bar charts with individual data points representing DAMs obtained from WT mice (red dots) and APP-KI mice (blue dots) aged 3, 6 and 9 months. Statistical significance was determined using unpaired Student’s t-test; \*\*\*\**P* < 0.0001.

**Fig. 2- figure supplement 1:**
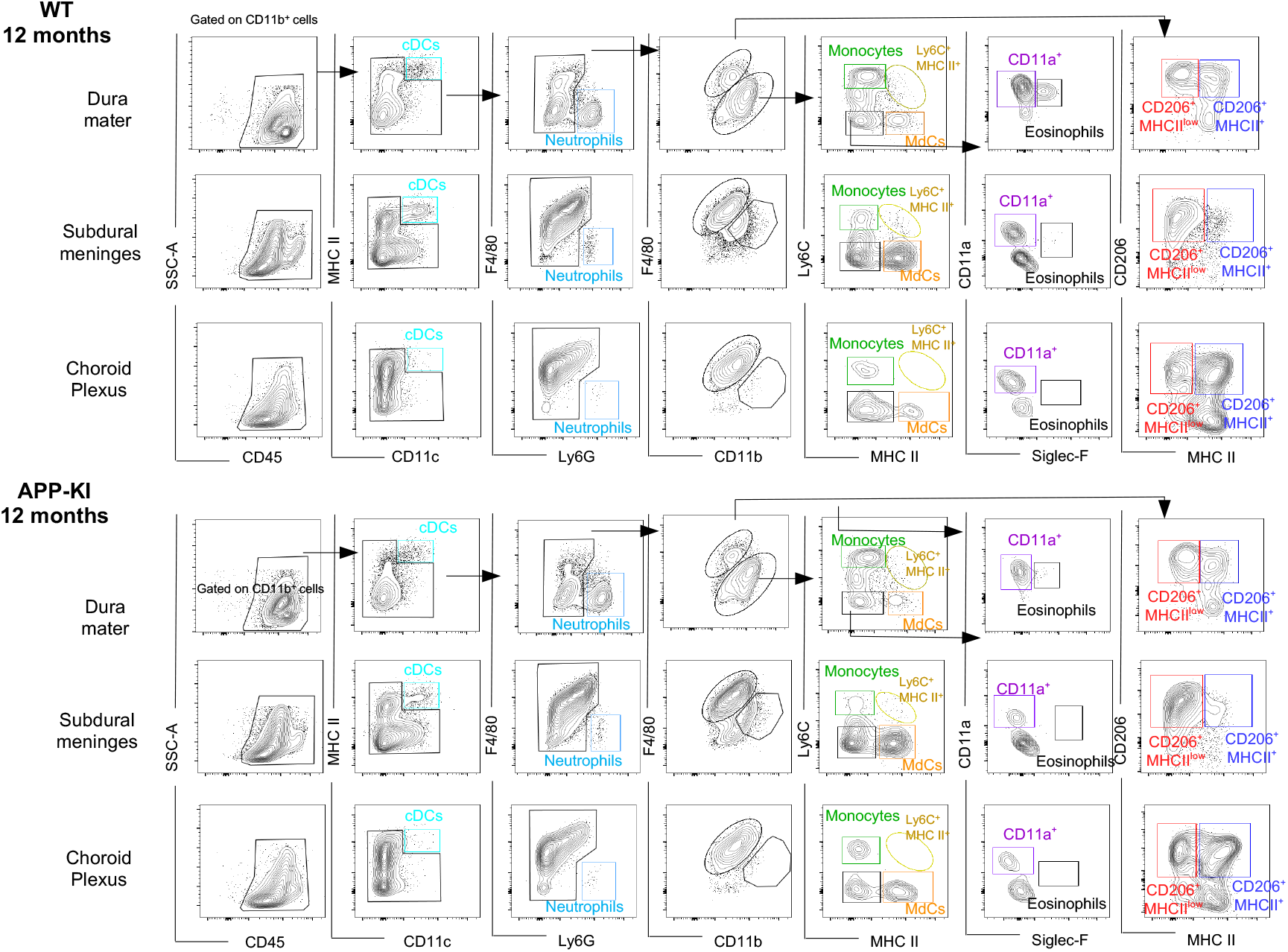
Characterization of myeloid cells in distinct brain border regions in health and AD. (A) Representative gating strategy used to visualize distinct myeloid cells subpopulations in the dura matter (DM), subdural meninges (SDM) and choroid plexus (CP) of WT and APP-KI mice aged 12 months. Cell classification as shown in legend for Figure 1.

